# Statistical framework for calling allelic imbalance in high-throughput sequencing data

**DOI:** 10.1101/2023.11.07.565968

**Authors:** Andrey Buyan, Georgy Meshcheryakov, Viacheslav Safronov, Sergey Abramov, Alexandr Boytsov, Vladimir Nozdrin, Eugene F. Baulin, Semyon Kolmykov, Jeff Vierstra, Fedor Kolpakov, Vsevolod J. Makeev, Ivan V. Kulakovskiy

## Abstract

High-throughput sequencing facilitates large-scale studies of gene regulation and allows tracing the associations of individual genomic variants with changes in gene expression. Compared to classic association studies, allelic imbalance at heterozygous variants captures the functional effects of the regulatory genome variation with smaller sample sizes and higher sensitivity. Yet, the identification of allele-specific events from allelic read counts remains non-trivial due to multiple sources of technical and biological variability, which induce data-dependent biases and overdispersion. Here we present MIXALIME, a novel computational framework for calling allele-specific events in diverse omics data with a repertoire of statistical models accounting for read mapping bias and copy-number variation. We benchmark MIXALIME against existing tools and demonstrate its practical usage by constructing an atlas of allele-specific chromatin accessibility, UDACHA, from thousands of available datasets obtained from diverse cell types.

**Availability:** https://github.com/autosome-ru/MixALime, https://udacha.autosome.org

## Introduction

Nowadays, genome- and transcriptome-wide association studies^1,2^ are the primary source of information on hereditary components of phenotypic traits, including disease susceptibility^3^. Yet, prioritization of causative genomic variants among multiple candidates remains a challenge, involving both statistical fine-mapping and functional annotation. The functional annotation of variant effects remains a no less challenge as the majority of disease-associated variants are non-coding^4,5^, and the diversity of mechanisms of gene regulation complicates the interrogation of non-coding variant effects.

Rich data facilitating analysis of non-coding variants is being obtained with modern omics technologies based on next-generation sequencing. Particularly, large-scale omics data facilitate the detection of heterozygous sites where the read counts are non-randomly distributed between homologous chromosomes. The allele-specific single-nucleotide variants and polymorphisms, AS-SNVs and AS-SNPs, are characterized by an imbalanced number of sequenced reads carrying one or the other allele^6^. Depending on the particular omics protocol, the allelic imbalance may reveal the association or direct variant impact on allele-specific transcription factor binding (AS-binding, ASB) using ChIP-Seq data^7–11^, allele-specific chromatin accessibility (ASA) using e.g. DNase-Seq or ATAC-Seq^8,12–16^, allele-specific DNA methylation^17,18^, and allele-specific gene expression per se^9,19–29^. From here on, we jointly call these observations allele-specific events (ASEs) related to sequence variant-dependent gene regulation.

The analysis of ASEs became a major topic at the crossroads of genetics and functional genomics, and various strategies for allelic bias identification have been developed in the past decade^30^. A workflow for the identification of ASEs starts with acquiring the list of SNVs, counting the reads supporting particular alleles at heterozygous sites, i.e., obtaining the SNV-level allelic coverage, and assessing the statistical significance of the observed allelic imbalance (see **Figure 1A**). While identifying candidate SNVs through variant calling stands on solid ground^31–33^, estimating the statistical significance of the allelic imbalance remains challenging.

**Figure 1.**
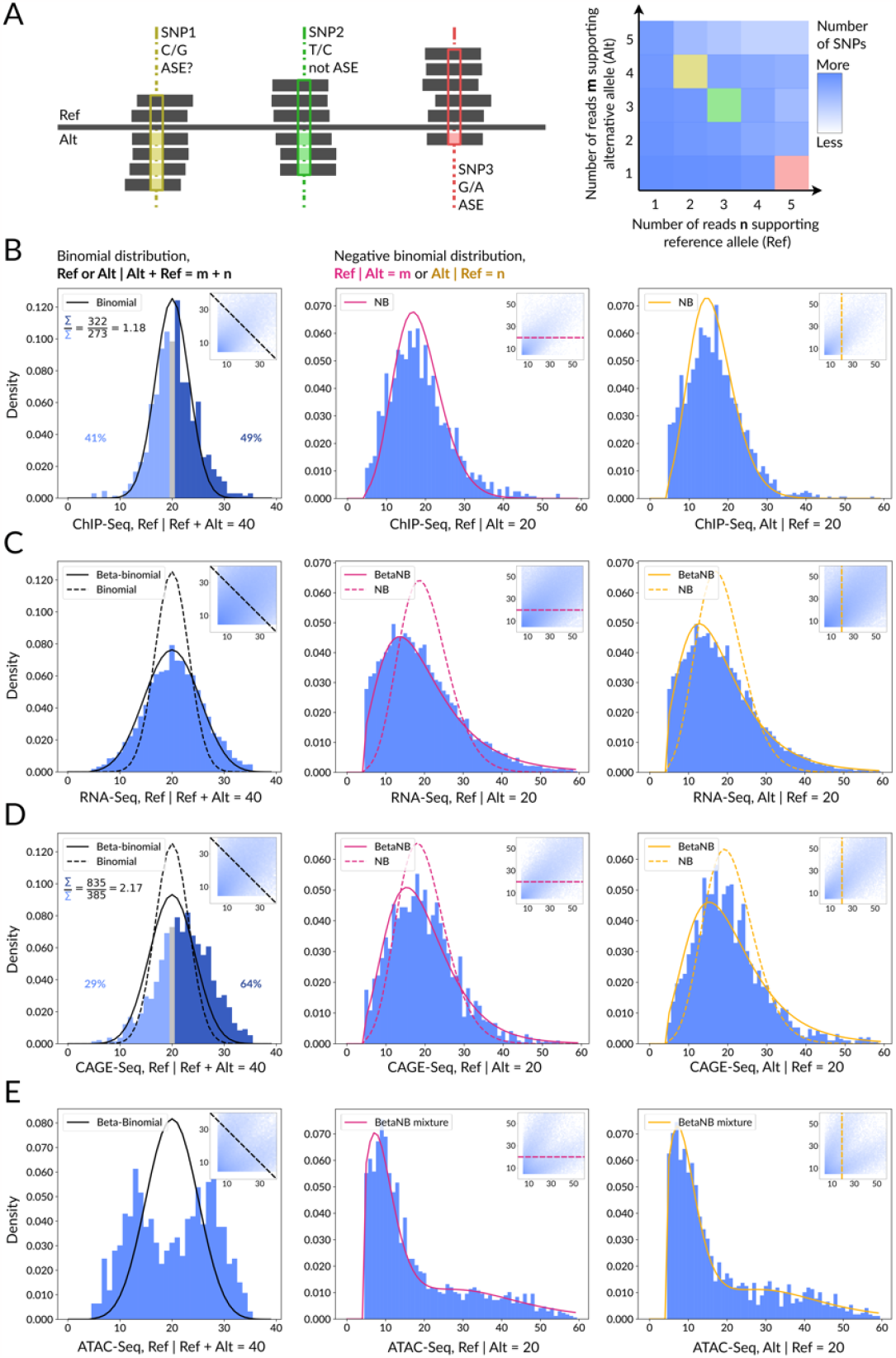
Challenges of ASE calling from various omics data resolved by different statistical models. **A:** A scheme representing the SNPs allele-specific coverages used as MIXALIME input is shown on the left. The total read counts distribution visualized as heatmap is shown on the right. **B:** An example of reference mapping bias found in 85 ChIP-Seq datasets from ADASTRA WA01 cells. The reference read counts distribution across the diagonal heatmap slice is shown on the left, fitted by the binomial model. The ratio of SNPs with the Ref counts > 20 (dark blue) to SNPs with the Ref counts < 20 (light blue) is shown in the top-left corner, percentages of both groups of SNPs are also shown on the sides of the histogram. The reference and alternative read counts distributions across the horizontal and vertical heatmap slices are shown on the right, fitted by the negative binomial model. **C:** An example of overdispersion found in 152 RNA-Seq datasets obtained from human left ventricles samples described in Sigurdsson et al. ^48^. The reference read counts distribution across the diagonal heatmap slice is shown on the left, fitted by the binomial (dash line) and beta-binomial (solid line) models. The reference and alternative read counts distributions across the horizontal and vertical heatmap slices are shown on the right, fitted by the negative binomial (dash line) and beta negative binomial (solid line) models. **D:** An example of extreme reference mapping bias and overdispersion found in 109 heart CAGE-Seq datasets from Deviatiiarov et al. ^53^. The histograms are plotted in the same manner as in (B) and (C). **E:** An example of CNVs with background allelic dosage, BAD, of 2 observed in ATAC-Seq datasets from UDACHA. The reference read counts distribution across the diagonal heatmap slice is shown on the left, fitted by the beta-binomial model. The reference and alternative read counts distributions across the horizontal and vertical heatmap slices are shown on the right, fitted by the mixture of beta negative binomial models. Abbreviations: ASE - Allele-Specific Event.

The source of the problem is the generally unknown expected distribution of allelic read counts in the case of allele balance, which is needed for statistical control. In some experimental setups, the control is available directly, for example, when studying asynchronous replication, where the pre-replication G1 cells were used to determine the unbiased expected ratio of allelic signals^34,35^. Yet, this is not the case for the majority of studies mapping transcription activity, DNA accessibility, or protein-DNA binding, which do not yield direct measurements of the allelic DNA dosage.

There are several technical sources of the excessive variability of the allelic read counts across the genome. First, the allelic imbalance is biased by heterogeneity at the level of cells or samples that inflates the read count variation. Second, false allelic imbalance may arise from genotyping errors, e.g., if somatic mutations present in a fraction of cells are mistaken for germline variants. The third major source is the reference mapping bias from the read mapping procedure in the absence of the personalized genome: the reads carrying non-reference alleles are either non-mapped or mismapped to wrong, sometimes multiple locations, with the tradeoff between dropout and mismapping rates controlled by the mismatch mapping penalty^36^. Consequently, depending on the read mapping strategy, the reads carrying the non-reference allele of a particular SNV become depleted on average, skewing both the overall and SNV-level allelic read counts. Without the personal genome at hand, this effect can only be countered with a variant-aware read realignment^36^.

Eventually, all these factors result in an overdispersed allelic count distribution that does not follow the convenient and commonly used binomial model where the allelic imbalance significance is assessed by a simple binomial test with the fixed success probability of *p* = ½^8,37–39^. A possible solution is to consider *p* as a random number coming from the beta distribution yielding the beta-binomial model^12,13,40–44^ (see **Figure 1B-D**, left subpanels). However, both the naive and beta-binomial approaches assume that the total number of reads at a genomic position is known, while in reality an unknown fraction of the total reads is missing, e.g., due to the read mapping bias.

An alternative model assumes that the number of reads supporting the reference or the alternative allele are measured independently and represent random variables following Poisson^45^ or negative binomial^46–50^ laws. The latter approach is similar to that used in differential gene expression tools like edgeR^51^. It is appropriate for haplotype-resolved data where the reads coming from paternal and maternal genomes can be fully separated, but less justified if the alleles and allelic read counts are not explicitly and unambiguously assigned to particular haplotypes or subgenomes. In this case, the convenient random variable can be the number of reads supporting the alternative allele given the fixed number of reads at the reference allele^10,52,53^ (see **Figure 1B-D**, right subpanels), resulting in the negative binomial model (see Methods for the substantiation).

In addition to the factors discussed above, the allele-specific analysis is complicated by the copy number variation (CNV)^42^. CNVs yield sporadic regions with increased allelic imbalance in normal cells, and, to a greater extent, in tumor samples and immortalized cell lines, which often exhibit global aneuploidy and other types of genome instability (see **Figure 1E**, left subpanel). Several solutions to counter this issue have been suggested. First, the CNV-affected regions can be tracked with the DNA input control. Many available packages for ASE calling rely on these data as a direct source of expected allelic read counts at the sites or regions of interest^13,28,37,39,40,54–65^. CNV-rich regions are then either excluded from consideration or processed in a special way, where an allelic copy number of 1:*L* is scored with a binomial test with *p* set to *1/(L+1)*^23,61,66^. Unfortunately, in many cases neither CNV profiling nor DNA control sequencing is available, or the latter has a shallow read coverage. Yet, it remains possible to account for the CNVs explicitly e.g. by estimating the background allelic dosage yielded by CNVs directly from the variant calls^10^. In the end, the allelic imbalance at variants in CNV regions is scored by the mixture model arising from the multiplied and non-multiplied alleles (see **Figure 1E**, right subpanel).

Here we present MIXALIME (MIXture models for ALlelic IMbalance Estimation), a versatile framework for identifying ASEs from different types of high-throughput sequencing data. We describe an end-to-end workflow from read alignments to statistically significant ASE calls, accounting for copy-number variation and read mapping biases. MIXALIME offers multiple scoring models, from the simplest binomial to the beta negative binomial mixture, can incorporate background allelic dosage, and account for read mapping bias. MIXALIME estimates the distribution parameters from the dataset itself, can be applied to sequencing experiments of various designs, and does not require dedicated control samples. We demonstrate MIXALIME performance against existing methods and present its large-scale application, UDACHA (Uniform Database of Allele-specific CHromatin Accessibility in the human genome), built by systematic ASE calling across 5858 human chromatin accessibility datasets.

## Results

### A general workflow from read mapping to ASE calling with MIXALIME

The overall workflow is shown in **Figure 2** and includes three stages: read mapping, SNP calling, and ASE identification. The read mapping can be performed by any modern software (e.g. BWA-MEM or STAR/HISAT2, depending on the data^67–69^). The SNP calling, e.g. with bcftools mpileup or GATK HaplotypeCaller^31,32^, must provide not only variant calls but allelic read counts. At the final step, MIXALIME comes into play: it fits the scoring model and estimates the significance of the allelic imbalance at each variant followed by obtaining combined *P*-values across individual samples or replicates and correction for multiple testing. This general scheme can be improved, first, by filtering the alignment with WASP^36^ after the SNP calling step to reduce the reference mapping bias in the absence of a personalized genome. Second, the genome-wide track of the relative background allelic dosage of aneuploid or CNV-rich samples can be estimated with BABACHI^10,70^. For commonly used immortalized cell lines this may significantly reduce the overdispersion and thus improve the reliability of ASE calling.

**Figure 2.**
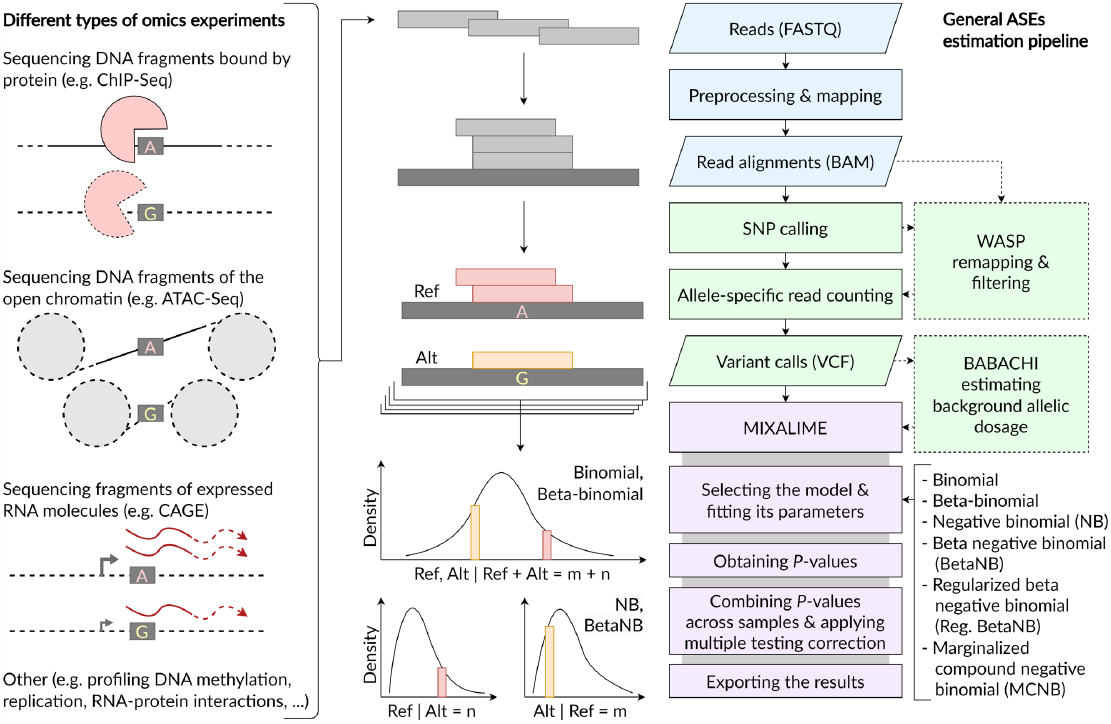
ASE calling with MIXALIME. Left: different types of omics experiments are suitable for calling allele-specific events, ASEs. Middle: schematic illustration of ASE calling steps: reads mapping, SNP calling, imbalance scoring. Right: ASE calling workflow. The reads remapping with WASP filtering, and background allelic dosage reconstruction with BABACHI are optional steps denoted with dotted lines. Abbreviations: ASE - Allele-Specific Event, SNP - Single Nucleotide Polymorphism.

### MIXALIME’s approach to ASE calling

The core idea of MIXALIME is the assumption that ASEs are rare events among multiple SNVs without significant allelic imbalance. Hence, the task can be reformulated as an outlier detection problem within a specified null distribution. The choice of the null distribution depends on the usage scenario and MIXALIME offers multiple models to handle allelic read count distributions with varying degrees of overdispersion. In addition to conventional binomial or beta-binomial distributions for allelic read count modeling, MIXALIME introduces means to explicitly address asymmetric mapping bias with left-truncated negative binomial (NB), beta negative binomial (BetaNB), and a novel marginalized compound negative binomial (MCNB) distributions, see Methods.

Let’s denote the read count at the allele 1 (e.g. the reference allele, Ref) as *x*, and the respective read count at the allele 2 (e.g. the alternative allele, Alt) as *y*. In MIXALIME we assume that the mean number of reads *r = E*[*y*] mapped to Alt across SNPs is linearly dependent on the respective read counts *x* mapped to Ref:

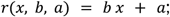

where are *a* and *b* are trainable parameters. Then, in the case of diploid CNV-free genome segments, *y* is assumed to be *F* distributed with a mean equal to *r*, where *F* is NB, BetaNB, or MCNB distribution, with the rate parameter *p* = ½ held fixed.

CNVs and aneuploidy increase the relative background allelic dosage, *BAD*, which is the local ratio of major to minor copy numbers^10^. In the case of *BAD* > 1, the read counts are modeled as a mixture of two *F* distributions with different fixed rate parameters 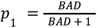 and *P*_2_ *=* 1 − *p*_1_. MIXALIME obtains maximum likelihood estimates (MLEs) of model parameters using gradient descent. In the end, SNVs are scored by computing *P*-values for a given *F* and its MLEs, and the *P*-values are combined across samples or replicates.

MIXALIME is implemented in Python with JAX framework for automatic differentiation and just-in-time compilation of the likelihood function, fitting the model is performed with scipy L-BFGS-B optimizer. MIXALIME can be installed with pip from PyPI repository, includes extensive built-in documentation, and accepts a common VCF format for input.

### MIXALIME performs ASE calling with adaptive sensitivity

To quantitatively assess the performance of different statistical models available in MIXALIME against existing state-of-the-art approaches, we used the cap analysis of gene expression data, CAGE-Seq, from 109 samples of 31 human hearts^53^. For MIXALIME, we followed the basic workflow (see **Figure 2** and Methods) skipping the estimation of background allelic dosage, as the samples from normal tissues should have diploid chromosome counts and CAGE yields a negligible coverage outside of transcription start sites thus not allowing to call CNVs or reconstruct BAD maps without external data.

For comparison, we performed ASE calling with existing tools that were suitable for the SNV-level CAGE data: QuASAR, originally designed for AS RNA-Seq, and BaalChIP, designed for detecting ASB in ChIP-Seq^40,42^. Both tools use the beta-binomial distribution to model read counts. For QuASAR, we combined resulting *P*-values across individuals in the same way as for MIXALIME allowing us to compare the results directly. Complete lists of significant ASEs from MIXALIME models, QuASAR, and BaalChIP can be found in **Supplementary Table S1**.

As there is no gold standard benchmark for ASEs, we estimated the reliability of ASE calls through ‘proxy’ by checking the overlap between the identified ASEs and known regulatory SNPs (rSNPs): (1) published ASBs from the ADASTRA database and (2) expression quantitative trait loci, eQTLs, from GTEx^10,71^. In brief, we assumed that a more sensitive method must detect more ASEs at a comparable significance threshold, whereas the method with a better specificity must provide a better overlap with ASBs from ADASTRA and eQTLs from GTEx. For a note, this definition of specificity and sensitivity has only an indirect relation to the respective classification performance metrics, as both eQTLs and ASBs encompass only subsets of true positive ASEs, most of which remain yet unidentified.

The beta-binomial model, implemented in QuASAR, demonstrates high specificity, i.e., yields a higher fraction of ASE calls overlapping with ASBs and eQTLs, but low sensitivity, i.e. generates the low number of detected ASEs in the whole range of significance thresholds (**Figure 3A**). However, at 0.05 FDR-corrected P-value (**Figure 3A, B**), MIXALIME’s beta negative binomial (BetaNB) model identifies three times more ASE counts than QuASAR (1676 as compared to 594 ASEs) with only a minor loss of specificity (the fraction of GTEx eQTLs drops from 0.49 to 0.46, while the ADASTRA ASB fraction remains at 0.55).

**Figure 3.**
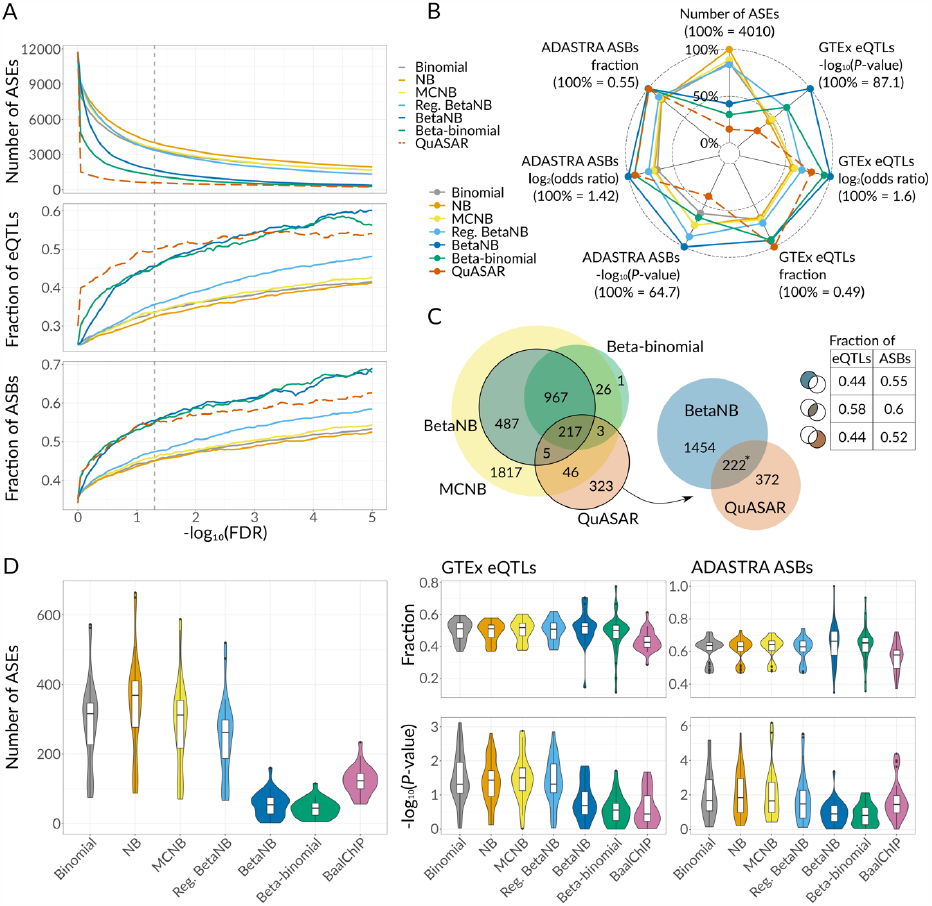
MIXALIME comparison to QuASAR and BaalChIP using heart CAGE data. **A:** The total number (top), the eQTLs-overlapping fraction (middle), and the ASB-overlapping fraction (bottom) of the significant allele-specific events (ASEs) detected at different FDR thresholds. The dashed line corresponds to QuASAR results. **B:** A radar plot demonstrating the number of significant ASEs (5% FDR), Fisher’s exact test results against eQTLs/ASBs (significance and odds ratios), and the respective fractions of ASEs. **C:** Left: Venn diagrams representing the intersection between ASEs called with three alternative MIXALIME models (beta-binomial, beta negative binomial, and marginalized compound negative binomial) and QuASAR, 5% FDR. Middle: the intersection between ASEs from MIXALIME BetaNB and QuASAR, the asterisk denotes the one-sided Fisher’s exact test *P*-value < 10^−15^, odds ratio > 10. Right: the fractions of eQTLs and ASBs among BetaNB-exclusive, QuASAR-exclusive, or overlapping ASEs. **D:** Violin and box plots demonstrating the number of ASEs (5% FDR) detected in each of 29 individuals, eQTL/ASB fractions, and Fisher’s exact test results for ASEs against the respective annotations. Abbreviations: ASB - Allele-Specific Binding event, ASE - Allele-Specific Event, eQTL - expression Quantitative Trait Locus, FDR - False Discovery Rate, MCNB - Marginalized Compound Negative Binomial, NB - Negative Binomial, Reg. BetaNB - regularized Beta Negative Binomial.

The alternative models including Binomial, NB, and MCNB, allow up to 8 fold increase in detected ASEs at 5% FDR at the cost of reduced overlap with known regulatory variants (e.g. for NB, eQTLs/ASBs fraction drops to 0.32/0.45, respectively). Finally, the Regularized BetaNB model yields intermediate results with 3357 ASEs of which 36% and 47% were annotated as eQTLs and ASBs, respectively. Thus, the desired priority of sensitivity over specificity and vice versa can be achieved with MIXALIME’s alternative scoring methods.

To estimate the significance of the observed overlap between ASEs and known rSNPs, we used Fisher’s exact tests, and the most conservative BetaNB model completely outperformed all other variants with Fisher’s log_2_(odds ratios) for eQTLs and ASBs of 1.6 and 1.4, and -log_10_(*P*-value) of 87 and 65, respectively.

Summing up, MIXALIME models offer multiple scenarios for ASE calling, including strict (BetaNB), balanced (BetaNB regularized at the model fit stage), or permissive (MCNB and NB) solutions with a desired trade-off between sensitivity and specificity (**Figure 3C**, left). Considering QuASAR, its ASEs overlap significantly the MIXALIME results with 222 ASEs shared with BetaNB (Fisher’s exact test *P*-value < 10^−15^, odds ratio = 10.4). The ASEs from the intersection expectedly show the highest fractions of eQTLs and ASBs, while 1454 BetaNB-exclusive cases have the same or even higher eQTL and ASB fractions compared to 372 cases exclusive to QuASAR (**Figure 3C**, right). Thus, compared to QuASAR, MIXALIME’s BetaNB reaches better sensitivity without loss of specificity.

Comparison to BaalChIP is a bit more complicated. First, by design, BaalChIP is applicable to groups of samples with a shared set of variants, e.g. belonging to one individual. Second, BaalChIP uses a Bayesian framework and provides not *P*-values but credible intervals of the allelic ratio estimates. Thus, for comparison against BaalChIP, MIXALIME was rerun with the *P*-values combined not across the whole set of samples but separately for each individual. In this setting, BetaNB and BetaBinomial MIXALIME models at 5% FDR performed similarly to BaalChIP in terms of the number of identified ASEs and ASB/eQTL overlap significance. However, the permissive models including Binomial, NB, MCNB, and Regularized BetaNB (**Figure 3D**), provided significantly better results as the statistical power in this comparison was limited by the number of samples and aggregation depth (2 to 6 samples) per individual.

Summing up, larger aggregation depth across more samples or replicates allows efficient application of strict BetaNB models, while permissive models suit better given a few samples or replicates, and in both scenarios, MIXALIME outperforms existing tools.

### MIXALIME efficiently accounts for mapping bias and CNVs

MIXALIME compensates for systematic mapping bias by fitting separate models for scoring the imbalance towards Ref and Alt alleles. However, it cannot account for site-specific bias, which can be handled by WASP filtering of the read alignment. To clarify the added value of WASP and also test MIXALIME performance at different sample sizes, we rerun the ASE calling with and without WASP using randomly selected 10, 25, 50, 75, or all 109 heart samples (see Methods and **Supplementary Table S2**). Each time we considered ASEs with 5% FDR, applied Fisher’s exact test against GTEx eQTLs, and estimated the relative P-value and odds ratio against the results of the Binomial scoring model as baseline (**Figure 4A**).

**Figure 4.**
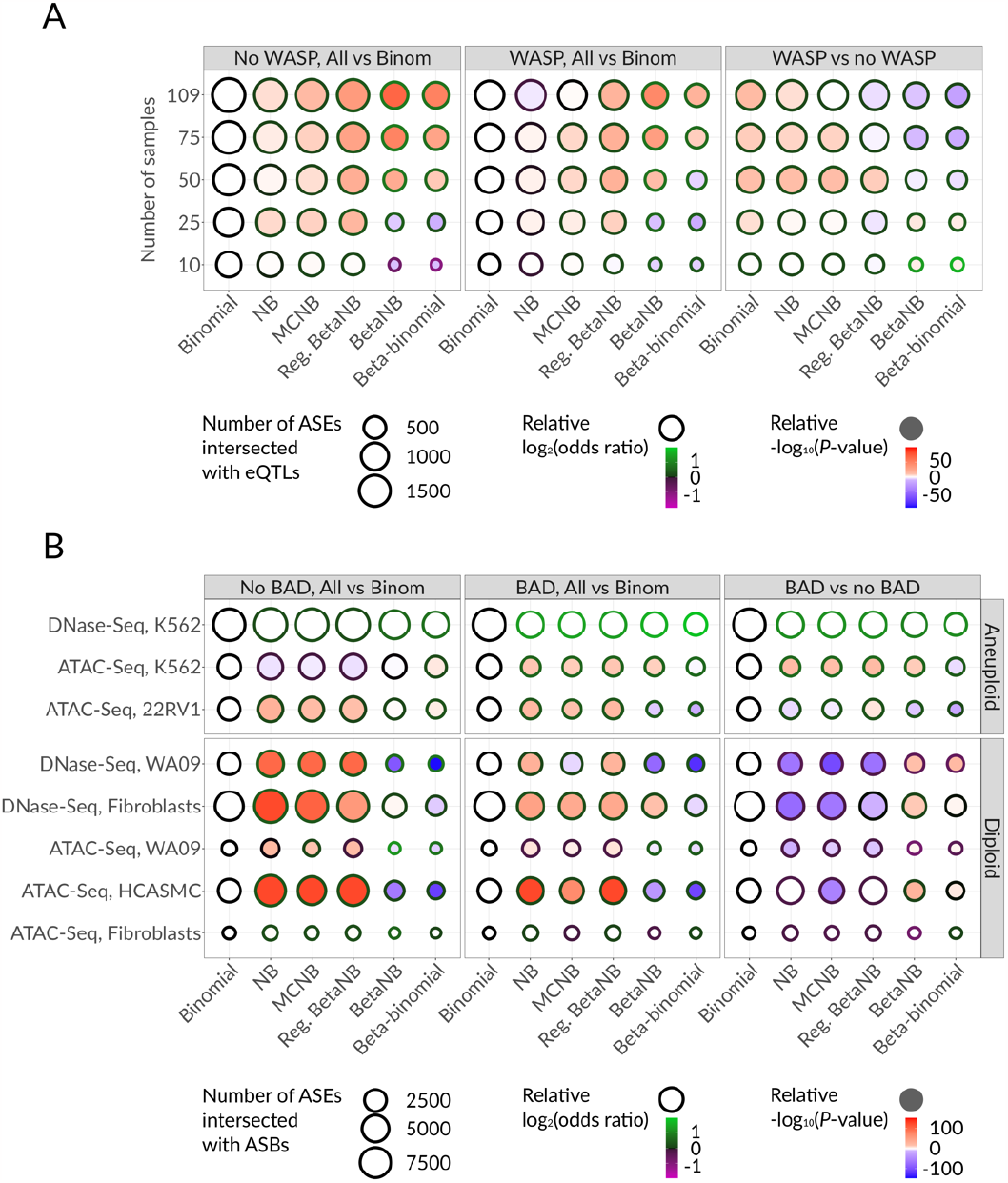
Performance assessment of MIXALIME’s alternative scoring models. **A:** Bubble plots illustrating the overlap between CAGE ASEs (with and without WASP filtering) and Heart eQTLs. The bubble size denotes the number of significant ASEs overlapping eQTLs, the fill and the border color denote the statistical significance and the log-odds ratio relative to the results of the Binomial model (left, middle) or relative to the results obtained without WASP (right), Fisher’s exact test. **B:** Bubble plots illustrating the overlap between ASEs and ASBs with and without accounting for the background allelic dosage, results are given relative to the Binomial model (left, middle) or relative to the no-BAD approach (right). Size, fill, and colors as in (A). Abbreviations: ASB - Allele-Specific Binding event, ASE - Allele-Specific Event, BAD - Background Allelic Dosage, eQTL - expression Quantitative Trait Locus, MCNB - Marginalized Compound Negative Binomial, NB - Negative Binomial, Reg. BetaNB - regularized Beta Negative Binomial.

As expected, in terms of agreement with eQTLs, stricter MIXALIME models perform better at higher sample sizes, both with and without WASP filtering. However, comparing WASP-filtered against unfiltered data, at large enough sample sizes WASP becomes unnecessary or even reduces the overall performance. Thus, as a rule of thumb, WASP filtering should be applied at small sample sizes as allowing to apply more permissive scoring models, but can be safely omitted otherwise.

It is also interesting to compare advanced MIXALIME models against the Binomial baseline. For example, when there are only 10-20 samples available, strict models including BetaBinomial and BetaNB show worse *P*-values compared to the Binomial model due to the limited number of ASEs, and it gets even worse without the WASP filter. On the other hand, when data from all 109 samples are aggregated, the BetaNB model demonstrates a two-fold increase in -log_10_(*P*-value) and log_2_(odds ratio) of overlap with eQTLs compared to those of the Binomial model (87.1 and 1.6 versus 40.8 and 0.9, respectively).

To estimate the effect of aneuploidy and CNVs on the ASE calling, we applied MIXALIME to 157 chromatin accessibility datasets (98 DNase-Seq and 59 ATAC-Seq), performed for presumably diploid (WA09, HCASMC, fibroblasts) and highly aneuploid (K562 and 22RV1) cells types (**Supplementary Table S3**). Each time, MIXALIME was used either with or without a BABACHI-generated BAD map, fitting a mixture or a basic model, respectively. We compared the detected ASEs with ASBs of ADASTRA, see **Figure 4B** and **Supplementary Table S4**. For chromatin accessibility data under study, permissive MIXALIME models achieve better performance in most cases. For aneuploid cells, BAD-aware mixture models are consistently more effective resulting in higher odds ratios for all tested cell lines and sometimes improving the significance as well. For normal diploid cells, allowing for BAD in fact reduces sensitivity, possibly due to clusters of neighboring AS sites being attributed to local CNV regions during BAD map reconstruction. For instance, the differences in -log_10_(*P*-value) and log_2_(odds ratio) between the BAD-aware mixture and basic models are 19.3 and 0.6, respectively, for regularized BetaNB applied to ATAC-Seq K562 samples, which are known to be mostly triploid, whereas ATAC-Seq performed on WA09 diploid cells results in -log_10_(*P*-value) and log_2_(odds ratio) differences equal -11.4 and -0.13, respectively. Finally, the BetaBinomial model scores poorly in all scenarios, suggesting its application should be avoided despite its popularity in AS studies.

### MIXALIME allows for differential ASE calling

In addition to basic ASE calling, MIXALIME allows the user to test differential allele specificity between two groups of samples. For this purpose, it starts by fitting the model parameters to the whole dataset, as in the standard ASE calling, with a fixed rate parameter *p*, e.g. with *p* = ½ in the diploid case. Next, for each SNP separately, MIXALIME frees the parameter *p* and fits it to the test and control groups separately obtaining the SNP-specific *p*_*test*_ and *p*_*control*_estimates. Finally, MIXALIME employs either Wald or likelihood-ratio test for evaluating whether the difference between *p*_*test*_ and *p*_*control*_is significant to identify differential ASEs. Conveniently, this approach is implemented for all model distributions present in MIXALIME.

For practical illustration, we reused the CAGE data for human hearts and split it by donor sex (53 male and 56 female samples). The two-group Wald test identified 73 sex-specific ASEs at 5% FDR of which 64 were present in the dbSNP common set (see **Supplementary Table S5**). Given the diversity of samples, it was possible to validate Ref/Alt ASE effects for particular SNPs by comparing the Alt/Alt versus Ref/Ref homozygotes. In our case, significant ASEs showed sex-specificity concordant with homozygous samples, while other variants did not demonstrate such tendency (Wilcoxon rank sum test *P*-value ∼0.05 for 23 sex-specific ASEs, *P*-value = 0.23 for other 1599 SNPs for which there were both hetero- and homozygous samples available, **Supplementary Figure S1A**).

Next, we used ANANASTRA^72^ to annotate these ASEs with ASBs, which resulted in 39 intersected SNPs (61%). Among transcription factors that bind these ASBs, ZFX, Zinc Finger X-Chromosomal Protein, turned out to be the most common with seven target ASBs, three of which (rs11730091, rs131804, rs28418438) were concordant with its binding motif. Two of those were annotated as GTEx eQTLs in the heart for numerous genes including *GATB* and *TYMP* involved in mitochondria functioning and angiogenesis. ZFX is differentially expressed between males and females (**Supplementary Figure S1B**, one-sided Wilcoxon rank sum test *P*-value < 10^−13^), suggesting a direct involvement of the respective ASEs in the sex-specific regulatory program of the human heart.

### Uniform Database of Allele-Specific Chromatin Accessibility

To showcase MIXALIME applicability to large-scale heterogeneous data, we performed ASE calling across the complete set of 5858 chromatin accessibility datasets uniformly reprocessed and available in GTRD (**Figure 5A, Supplementary Table S6**)^72,73^. SNP calling was performed with GATK using read alignments from 1850/3801/207 DNase-/ATAC-/FAIRE-Seq experiments; we kept only sufficiently covered common heterozygous SNPs (0/1 in VCF GT field). The next steps included genotype- and metadata-based sample clusterization and subsequent BABACHI background allelic dosage estimation run separately for each group of related samples sharing cell types and variant calls (see Methods). Finally, we applied MIXALIME and detected 142312/105901/1167 total allele-specific chromatin accessibility sites at 84028/56014/1162 rsSNPs across 240/359/10 cell types, respectively. Most of the ASEs were unique for DNase-Seq or ATAC-Seq data even at the level of rsSNP IDs (**Supplementary Figure S2A**), primarily due to different sets of assessed cell types. Yet, for intersecting significant ASEs passing 5% FDR in matched DNase- and ATAC-Seq cell types, the allelic imbalance was mostly concordant, likely revealing the true effect these ASEs reflect or induce in the corresponding chromatin regions (**Supplementary Figure S2B**).

**Figure 5.**
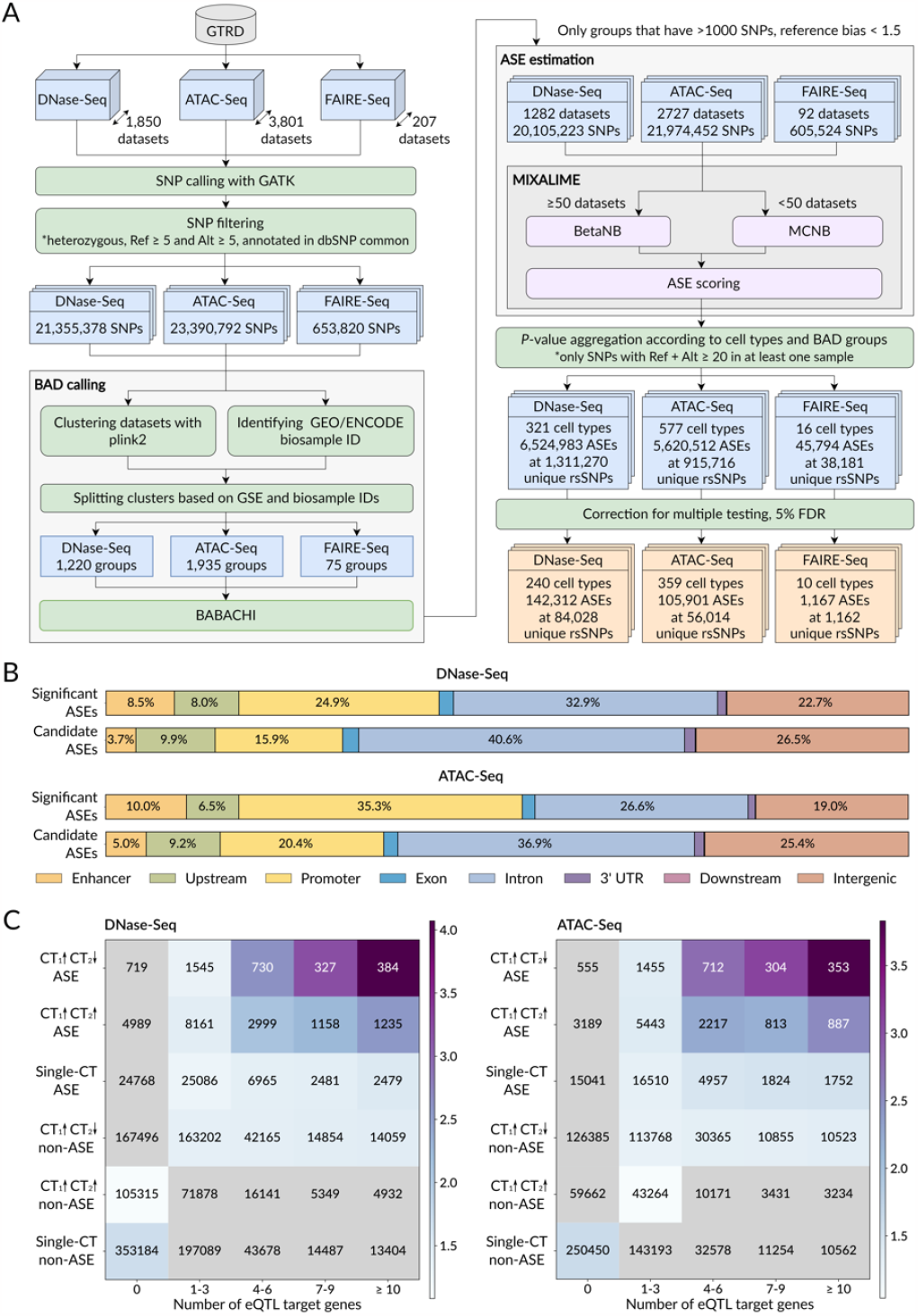
UDACHA, a uniform large-scale database of allele-specific chromatin accessibility in the human genome. **A:** UDACHA workflow. First, SNP-calling with GATK is performed from GTRD read alignments. Second, samples are grouped according to cell types and plink2 clusters, followed by BABACHI group-specific BAD reconstruction. Finally, MIXALIME is used to call ASEs. In the end, at 5% FDR we identified 142,312 / 105,901 / 1,167 ASEs in DNase-Seq / ATAC-Seq / FAIRE-Seq data, respectively. **B:** Chromatin ASEs distribution among different types of genomic regions. The complete bars correspond to the full sets of rsSNPs coinciding with ASEs called from DNase-Seq (top) and ATAC-Seq (bottom). Compared to candidate variants, the significant ASEs are more often found in promoters and enhancers. The percentage of ASEs falling into particular types of genomic regions is shown on bar labels. The top bar in each subpanel: significant ASEs passing 5% FDR, 84028 sites for DNase-Seq and 56014 sites for ATAC-Seq; bottom bar: candidate ASEs passing the coverage thresholds and tested for significance, 1227242 sites for DNase-Seq and 859702 for ATAC-Seq. The annotation procedure is the same as in ADASTRA^10^. **C:** eQTL Enrichment of DNase-Seq (left) and ATAC-Seq (right) rsSNPs of different categories. Y-axis: CT_1_↑CT_2_↓, rsSNPs with ASEs but opposite allelic preferences in different cell types; CT_1_↑CT_2_↑, rsSNPs with ASEs and same allelic preferences across cell types; Single-CT, ASEs significant in a single cell type. Non-ASEs: SNVs with FDR above 0.05. X-axis: the number of eQTL target genes according to GTEx eQTL data. The coloring denotes the odds ratios of the one-tailed Fisher’s exact test for a particular cell of the table against all other rsSNPs in the table. The gray cells correspond to non-significant enrichments with P > 0.05 after Bonferroni correction for the total number of tested cells. The values in the cells denote the numbers of rsSNPs. Abbreviations: ASE - Allele-Specific Event, BAD - Background Allelic Dosage, BetaNB - Beta Negative Binomial, CT - Cell Type, eQTL - expression Quantitative Trait Locus, FDR - False Discovery Rate, MCNB - Marginalized Compound Negative Binomial, SNP - Single Nucleotide Polymorphism, UTR - UnTranslated Region.

To further verify the reliability of ASE calls for selected cell types, we compared the respective open chromatin ASEs against ADASTRA ASBs for 6 cell lines with multiple candidate ASBs in ADASTRA and candidate ASEs in UDACHA and found the statistically significant overlap of ASEs with ASBs as well as a correlated allelic imbalance (**Supplementary Table S7**). However, the absolute numbers of coinciding SNPs were not high. Thus, UDACHA assesses quite a different quantity than ADASTRA highlighting allelic imbalance at many previously unexplored rSNPs even in well-studied cell types such as K562 or MCF7. We also annotated the genomic localization of UDACHA ASEs (**Figure 5B**, compare with **Figure 4C** in ADASTRA^10^). As expected, significant ASEs compared to all candidate sites prefer promoter and enhancer regions, although, in absolute counts, introns and intergenic regions carry many ASEs as well.

Finally, we used GTEx to estimate whether different groups of SNPs displayed concordant or discordant allelic imbalance between cell types are special in terms of the eQTL-reflected association with gene expression. As in ADASTRA, the switching ASEs with ‘antiparallel’ behavior, i.e. with opposite allelic preference in different cell types, affected more target genes according to GTEx, both for DNase- and ATAC-Seq ASEs, suggesting such variants affect promiscuous enhancers regulating transcription of multiple genes in cell type-specific manner (**Figure 5C**).

## Discussion

ASE calling methods are diverse and rapidly evolving. There are methods that avoid the classic binomial or beta-binomial approach but nonetheless share their limitations, such as using the chi-square goodness of fit test^19,74^ or G-test^56^. In special cases, such as identifying allele-specific topologically associating domains^75^ or allele-specific DNA replication and methylation, the authors compared Z-scores against a fixed threshold, performed Fisher’s exact test, or chi-square test of independence with categorical variables e.g. such as allele (Ref/Alt), the molecule type (RNA/DNA), the cell cycle phase (G1/S), or cytosine methylation state (methylated/unmethylated)^18,39,76,77^. Yet, with so many methods for ASE calling, only a few are generally applicable to call ASEs from a generic set of heterozygous SNVs and do not rely on control DNA sequencing, phased haplotypes, or extensive replication. Further, the majority of existing packages are designed with RNA-Seq data in mind and estimate gene-, transcript- or exon-level allelic imbalance taking into account that closely located SNVs in mRNAs can physically share the allele-specific expression. A simple approach is to leave a single top SNV per feature, e.g. with the highest read count^27,28,40^, but there are better strategies based on merging allele-specific counts on individual SNVs through phasing^37,62,65^, pseudo-phasing^58,61,63^, or test statistics combination^60^, thereby improving but complicating the pipeline and limiting applicability to the other types of sequencing data.

In this study, we described MIXALIME, a generally applicable ASE calling software. MIXALIME can handle the standard VCF files and score the SNV-level allelic imbalance with multiple models using the allelic coverage at individual SNVs. MIXALIME allows the user to combine the results across any number of samples and takes into account the background allelic dosage induced by CNVs. In contrast to many other tools, it does not require phased genomes, DNA input control, or some specific experimental design that includes a large number of replicates. The default MIXALIME pipeline allows direct ASE estimation from VCF files yielded by an SNP-caller (**Supplementary Figure S3**).

MIXALIME is user-friendly and offers a number of options for fine-tuning ASE calling thus enhancing its applicability. First of all, the user may use WASP to correct the mapping bias that can affect subsequent ASE estimation especially when the number of samples is small (**Figure 4A**), but it is not mandatory. Second, CNVs can be taken into account if the samples are assumed to be polyploid, which can use either the externally obtained CNV profile or a BAD map reconstructed with BABACHI (**Figure 4B**). Finally, depending on the size of the input data, the optimal ASE scoring model may vary from strict BetaNB to permissive NB and MCNB, with the compromise regularized BetaNB model being applicable in most cases.

From our test with 109 CAGE experiments, we came up with a general recommendation to run BetaNB when the number of samples exceeds 50, MCNB when the number of samples is 10 or less, and regularized BetaNB otherwise (**Supplementary Figure S3**). While we cannot guarantee that these recommendations would hold for any experiment type with arbitrary coverage, MIXALIME provides detailed model fit plots that can be used to guide the selection of the model and ensure that the mode distribution adequately fits the observed data (**Supplementary Figure S4**).

Overall, MIXALIME allows the user to estimate allele-specific events using different types of experiments across hundreds of samples in a relatively short time compared to the other existing ASE calling tools. In particular, for MIXALIME it takes about 15 minutes to estimate ASE significance for 109 heart samples starting from VCF files and using 1 modern computing core, which is more than nearly half a minute used by QuASAR but remains negligible compared to the computational load of read mapping and SNP calling. Further, MIXALIME offers multithreading out-of-the-box, does not require extra data pre-processing, and works significantly faster than BaalChIP which takes more than 9 hours to identify ASEs.

Taking advantage of MIXALIME versatility, we created an allele-specific chromatin accessibility database UDACHA from 5858 uniformly reprocessed chromatin accessibility datasets which included 1850 DNase-Seq, 3801 ATAC-Seq, and 207 FAIRE-Seq experiments available in GTRD. The resulting database includes 249380 ASEs in total that are preferably located within promoter and enhancer regions compared to other candidate SNPs and enriched by GTEx eQTLs (**Figure 4B, C**) and ADASTRA ASBs (**Supplementary Table S7**). In addition to the variants that have been already annotated in other databases, UDACHA includes many newly discovered rSNPs and makes use of different experiment types that complement each other.

To sum up, in this paper we described MIXALIME, the versatile user-friendly tool for ASE estimation using various data types, and UDACHA, the comprehensive allele-specific chromatin accessibility database. We believe these resources will help the scientific community decipher gene regulation and facilitate studies of regulatory variants by turning the complicated analysis of omics allele-specificity into a routine, reliable, and easily reproducible procedure.

## Methods

### MIXALIME models

Given allelic read count data, MIXALIME can fit several statistical models (see below) under the following assumptions:

1. ASEs are rare, hence their identification can be framed as an outlier detection problem;
2. The mean *r* of the read count at the allele 2 (e.g. the alternative allele, Alt) linearly depends on the read count *x* at the allele 1 (e.g. the reference allele, Ref), and vice-versa:

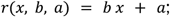
3. Read counts at an allele 2 (or allele 1) are distributed according to a law *F* with a probability density function *f*.

Specifically, *F* can be negative binomial (NB), beta negative binomial (BetaNB), BetaNB, or marginalized compound negative binomial (MCNB). The NB-based distributions have an advantage of over binomial distribution as they allow for the assumption 2 that the mean number of reads mapped to one allele (proportional to *r*) is linearly dependent on the read counts mapped to the other allele. A brief overview of models is given below, see Supplementary Methods^78^ for details and a formal substantiation. MIXALIME uses the left-truncated at *l* variant of distributions, as SNVs with low coverage at a particular allele should be pre-filtered to avoid technical false positive variant calls and somatic mutations. In this study, we use *l* = 4, i.e. only counts ≥ 5 are allowed. The general MIXALIME model is

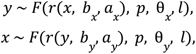

where θ is an optional extra parameter controlling for variance (e.g., κ, see BetaNB below). Note that estimatable parameters of the model are considered independently for the distributions of reference allele count *x* and alternative allele count *y*. In each case, we assume the other allele to be fixed (e.g. for the distribution of the read counts *x* at the reference allele, we assume the read count *y* at the alternative allele to be known). The key feature of this approach is that it enables separate scoring of allelic imbalance favoring each of the two alleles. This way we model the reference mapping bias implicitly as a difference between *r* parameters for reference and alternative distributions.

Once *F* is chosen, the model parameters are estimated with either the maximum likelihood (ML) or, for regularized BetaNB, with a maximum-a-posteriori approach, see Section 7 in Supplementary Methods^78^. MIXALIME provides the following set of models.

#### NB

To use this model, we assume a linear read mapping bias and a negative binomial distribution of *y* for a given *x*, and, symmetrically, the same for *y*:

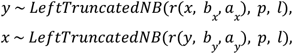

where *LeftTruncatedNB*, is the left truncated at *l* negative binomial distribution, *r* can be considered to reflect an expected number of read counts at a fixed allele, *b*_*x*_, *b*_*y*_, *a*_*x*_, *a*_*y*_ are active parameters to be estimated, *p* is a fixed rate parameter equal to 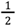 in case of diploid and CNV-free genomic regions.

#### BetaNB

Similarly to the generalization from a binomial to a beta-binomial model, we can assume *p* ∼ *Beta*(*α, β*) and apply a convenient reparametrization of *Beta* in terms of its mean μ and “concentration” κ^79^:

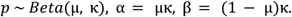

This compound model can be marginalized to the beta negative binomial distribution by integrating out *p*, therefore:

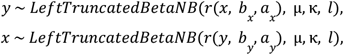

where *LeftTruncatedBetaNB* is the left truncated at *l* beta negative binomial distribution.

#### Regularized BetaNB

BetaNB model provides conservative *P*-value estimates as the beta negative binomial distribution at small values of κ is very heavy-tailed compared to the negative binomial distribution, and small κ may provide the optimal fit for the datasets with lots of ASEs consequently penalizing their significance. Thus, on the one hand, it is convenient to be able to compromise the goodness of fit for greater sensitivity by encouraging higher values of κ. On the other hand, high coverage data has lower variance, i.e. higher κ are expected at higher values of the fixed allele read count *c*. We introduce a regularization that accommodates this observation by assuming that the reciprocal of κ follows Laplace distribution:

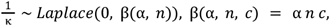

where β is a scale parameter, α is a regularization multiplier/hyperparameter, and *n* is a total number of observations. Here, the scale parameter gradually increases as we slide farther across the dataset towards higher coverage at a fixed allele, and the size multiplier *n* makes α more dataset-agnostic.

#### MCNB

The NB and BetaNB models are built on the assumption that the read count at the fixed allele is known. In practice, the read count at the preselected allele is not measured exactly and should be considered a random variable itself. Let’s assume that the alternative allele read count *y* is distributed as a zero-truncated binomial random variable (*ZTBin*). The zero-truncation is necessary to accommodate for the two facts: the allele-specificity is undefined for homozygous SNVs, and, technically, *r* > 0 in NB.

Let’s consider the following model:

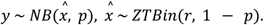

It turns out that 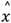 can be marginalized out and a marginal distribution of *y* can be obtained (see the proof in Appendix C of Supplementary Methods^78^):

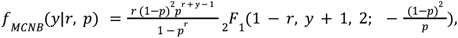

where _2_*F*_1_ is the Gauss hypergeometric function.

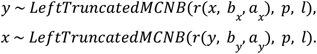

Thus, the resulting MCNB model is

#### Mixture models

Suppose that 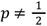 (or 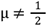 in the case of BetaNB), which happens for SNVs located in CNVs or duplicated chromosomes. For instance, there might be three copies of a maternal allele and one copy of a paternal allele in a tetraploid organism, which results in 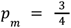 and 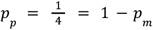. Most of the time, the completely phased personal genome and even partial haplotypes are not available, i.e. the exact number of copies of the reference and the alternative allele for any particular SNP remain unknown. However, it remains possible to estimate the ratio of the major to the minor allele copy numbers, that is the relative background allelic dosage (BAD), directly from SNP calls with an unsupervised approach or from an experimentally obtained CNV map^10^.

We tackle this problem by assuming that each read is coming from one (e.g. maternal) chromosome with probability *w* and from the other chromosome (e.g. paternal) with probability 1 − *w*, where BAD defines the balance between *w* and 1 − *w*. This is done naturally with the mixture distribution:

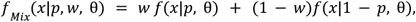

where *f* is either NB, BetaNB, or MCNB distribution function, θ is a parameter vector with *w* and *p* excluded, 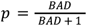, and *w* is a weight in the mixture model, an active parameter to be estimated from the data.

#### Legacy models

For convenience and benchmarking, MIXALIME provides conventional binomial and beta-binomial models. Note that those models do not use the mixture distributions to model count data for 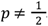. However, for the beta-binomial model, estimate the concentration parameter κ is separately estimated for each BAD.

#### Fitting the models to the observed read counts

Line *r* parameters *a, b*, and, when applicable, mixture and concentration parameters *w* and κ can be estimated with ML. However, to accommodate for possible non-linearities in the behavior of *r* depending on the fixed allele, we use local ML^81^ instead: a dataset is separated into a set of overlapping windows of a constant size centered at each fixed allele read count, and ML estimation is done for each window separately (see Section 7.3 of Supplementary Methods^78^ for more details).

### MIXALIME workflow and implementation

MIXALIME usage implies going through several steps (see **Supplementary Figure S3** and **Figure 5** in Supplementary Methods^78^), including model parameter estimation (fit), calculating raw *P*-values for each unique pair of read counts (reference allele count *x*, alternative allele count *y*), combining raw *P*-values across user-defined groups of samples, correcting the *P*-values for multiple testing, and, finally, exporting results in a tabular form. Optionally, it is possible to inspect the goodness of fit and, if necessary, identify the differential ASEs between two groups of samples.

MIXALIME accepts variant calls in either VCF or BED-like format, the BAD maps can be provided either directly in BED-like files or, for VCF, as separate files. Once MIXALIME is done preprocessing input data, it packs all the necessary information into a single project file. This enables MIXALIME projects to be easily portable between machines without keeping the initial variant files.

Once the model parameter estimates are obtained, ASEs can be identified in individual samples, and right-tailed *P*-values across samples of user-defined groups are combined with the Mudholkar-George *logitp* method^82^. For individual samples, the ASE effect size is estimated as 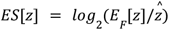, where *z* is either the reference allele read count *x* or the alternative allele read count *y*, and 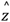 is the observed read count. The combined ASE effect size is estimated across samples as a weighted average using − *log*_10_*P*-values as weights. For technical details of computing p-values for various distributions, see Section 9.1 in Supplementary Methods^78^.

MIXALIME is implemented in Python 3.7 programming language. It relies on JAX framework for automatic differentiation of the log-likelihood function and its just-in-time compilation for optimal performance. The optimization of the ML problem is performed with the L-BFGS-B routine provided by the scipy package. In the process, the parameter *w* is bounded in interval (0, 1), the concentration parameter κ is bounded in interval (0, 10000) with the upper bound necessary to make the optimizer avoid attempting to rise κ to infinity if applied to underdispersed datasets. MIXALIME parameter estimation procedure is parallelized across BADs, and the choice of a fixed allele, and the scoring procedure is parallelized across SNVs. Computation of p-values is done with the help of arbitrary precision algebra package gmpy2, which proved to be crucial for computing CDFs of BetaNB and MCNB distributions. A user interacts with MIXALIME using a command-line interface, with each step (reading/preprocessing input data, estimating parameters, computing *P*-values, combining *P*-values across groups, exporting data) invoked with a separate command. The documentation is accessible using the `--help` command line argument.

### Overview of the sequencing data used in the study

In this study we used several high-throughput sequencing datasets obtained from different sources.

1. To illustrate the distributions of allele-specific read counts we used 85 ChIP-Seq datasets obtained WA01 cells and processed in ADASTRA^10^ (used in **Figure 1B)**, 152 human heart left ventricle RNA-Seq samples from Sigurdsson et al.^48^ (used in **Figure 1C**), 109 human heart CAGE-Seq samples from Deviatiiarov et al.^53^ (used in **Figure 1D**), and 59 ATAC-Seq datasets of UDACHA listed in **Supplementary Table S3** (used in **Figure 1E**);
2. 109 human heart CAGE-Seq samples from Deviatiiarov et al.^53^ were used to compare MIXALIME with QuASAR (**Figure 3A-C** and **Figure 4A**), two ‘singleton’ samples were excluded from the comparison against BaalChIP (see **Figure 3D**), see below for details; the same 109 CAGE-Seq samples were used in analysis of the sex-specific differential allelic imbalance (**Figure S1**);
3. To evaluate the effect of CNVs on the naive and BAD-aware allele-specific analyses (**Figure 4B**), 59 ATAC-Seq and 98 DNase-Seq samples of UDACHA listed in **Supplementary Table S3** were used;
4. 3801 ATAC-Seq, 1850 DNase-Seq and 207 FAIRE-Seq samples (**Supplementary Table S6**) from GTRD^73^ were used to build the UDACHA database.

### CAGE data and preprocessing pipeline

In the primary analysis, we used 109 FASTQ files of CAGE-Seq experiments performed on 31 human hearts and described in ^53^. The reads were mapped against hg38 human genome using hisat2 (v.2.2.1) with comprehensive GENCODE annotation (v.39) and `--very-sensitive` preset. Second, *filter_reads*.*py* Python script (Stamatoyannopoulos Lab github) was used to filter out the reads containing more than 2 mismatches and mapping quality < 10. Next, SNP-calling and allelic read counting were performed using the approach of ^12^ for DNase I processing pipeline and included the following steps: (1) bam files from different samples of one individual were merged using samtools (v.1.10) `merge`; (2) SNP-calling was performed using bcftools (v.1.10.2) `mpileup` with `--redo-BAQ --adjust-MQ 50 --gap-frac 0.05 --max-depth 10000` and `call` with `--keep-alts --multiallelic-caller`; (3) the resulting SNPs were split into biallelic records using bcftools `norm` with `--check-ref x -m -` followed by filtering with bcftools `filter -i “QUAL>=10 & FORMAT/GQ>=20 &FORMAT/DP>=10” --SnpGap 3 --IndelGap 10` and bcftools `view -m2 -M2 -v snps` leaving only biallelic SNPs covered by 10 or more reads; (4) SNPs were annotated using bcftools `annotate` with `--columns ID,CAF,TOPMED` and dbSNP (v.151) ^83^, heterozygous variants located on the reference chromosomes with GQ ≥ 20, depth ≥ 10, and allelic counts ≥ 5 for each allele were filtered for each individual with awk (v.5.0.1), (5) optionally, WASP (v.0.3.4) was used together with hisat2 and `filter_reads.py` to find, remap, and filter the SNP-overlappind reads that failed to map back to the same location after the alleles swapping according to WASP procedure, (6) Python `count_tags_pileup_new.py` script was used to perform sample-level allelic read counts with pysam (v.0.20.0) and `recode_vcf.py` was used to convert the resulting BED files to VCF. Additionally, to produce VCF files containing both heterozygous and homozygous positions, the heterozygous filter from step (4) was turned off.

To estimate MIXALIME models performance using a different number of heart samples, i.e. 10, 25, 50, and 75 randomly chosen samples out of 109, ASE estimation procedure was repeated 5 times for each number of samples, mean Fisher’s exact test -log_10_(*P*-values), log_2_(odds ratios) and number of ASEs are shown in Figure 4A with the full results available in **Supplementary Table S2**.

### Application of existing ASE calling methods

To assess MIXALIME performance against other available instruments, we performed ASE calling with QuASAR (v.0.1) and BaalChIP (v.1.0.0).

ASE calling with QuASAR was performed from WASP-filtered input VCF files with homozygous positions and minor allele frequencies (MAF) from 1000 genomes as listed in dbSNP (v.151). According to the QuASAR manual (https://github.com/piquelab/QuASAR), we estimated the priors for the genotypes from MAF and used `fitAseNullMulti` for each individual separately to perform genotyping followed by the ASE inference with `aseInference`. To obtain the resulting significance estimations, we combined the sample-level *P*-values via *logitp* function from the metap R package (v.1.8) and performed Benjamini-Hochberg multiple testing correction as in MIXALIME.

To identify ASEs with BaalChIP, the BAM files obtained after WASP filtering were used together with the TSV files containing the per-individual heterozygous positions and the BED file containing the full-length chromosome regions. These data were passed to BaalChIP and `alleleCounts` were used with default parameters to perform allele-specific read counting as suggested by the BaalChIP authors (https://github.com/InesdeSantiago/BaalChIP). Next, `mergePerGroup`, `filter1allele`, and `getASB` were used to identify allele-specific events. The single-sample data (no replicates) from two individuals were excluded before the read-counting leaving 107 samples from 29 individuals. To make a fair comparison, MIXALIME results were re-collected for the same set of samples.

### Chromatin accessibility data and UDACHA pipeline

UDACHA is built from short read alignments produced with bowtie2^84^ against hg38 genome assembly which were stored internally in the GTRD database^73^. The UDACHA pipeline was mainly inherited from ADASTRA^10^. Briefly, PICARD was used for deduplication, followed by GATK base quality recalibration and variant calling with GATK HaplotypeCaller. The dbSNP 151 common variant set was used for annotation^83^. The resulting variant calls were filtered to meet the following requirements: (1) an SNV must be biallelic and heterozygous (GATK annotation GT = 0/1); (2) have read coverage ≥ 5 at both the reference and alternative alleles; (3) listed as an SNP in the dbSNP 151 common set.

To improve the reliability of BAD calling, the samples were segregated into BAD groups of the same cell type, experiment series (by GEO GSE or ENCODE biosample), and sharing a similar set of variants. The latter was checked by running plink2^85^ (`plink2 --allow-extra-chr --threads 20 --make-king square`), zeroing inter-sample distances < 0.4, and clustering the samples (complete linkage, correlation metric) at the 0.1 distance threshold. For each BAD group of samples, BABACHI^70^ [ZENODO doi: 10.5281/zenodo.7901610] was executed with `--states “1,4/3,3/2,2,5/2,3,4,5,6” -p geometric -g 0.98`. Samples with extreme mapping bias N_(Ref>Alt)_ / N_(Ref<Alt)_ > 1.5 were excluded from each BAD group, here N_(Ref>Alt)_ is the number of variants with higher read count at the reference allele. The BAD groups with less than 1000 variant calls in total were excluded.

Finally, separately for DNase-Seq, ATAC-Seq, and FAIRE-Seq, we have fitted the MIXALIME MCNB and BetaNB models. For ASE calling, MCNB was used as a default permissive model, and BetaNB was used for the cell types with more than 50 samples.

### Annotating UDACHA ASEs with genomic locations and eQTLs

The annotation procedure for **Figure 5B** was inherited from ADASTRA. To analyze an overlap between ASEs of different types and eQTLs (**Figure 5C**), we used all significant “variant, gene” pairs from GTEx (release V8)^71^.

## Data Availability

MIXALIME is available at pypi, github and ZENODO [doi:10.5281/zenodo.8146508]. UDACHA database is freely accessible at https://udacha.autosome.org and the source code of the pipeline is available at github and ZENODO [doi:10.5281/zenodo.8191902].

## Acknowledgments

The study was primarily supported by RSF [20-74-10075 to I.V.K.]. Analysis of human heart CAGE was supported by the Ministry of Science and Higher Education of the Russian Federation (grant no. 075-152-021-601). We thank Daria Bykova for her help with the genomic annotation of UDACHA ASEs. We thank Niagara Computers LLC and personally Valery Egorshev for valuable technical support in setting up computational resources. We wholeheartedly thank Prof. Sigurdsson for sharing the precomputed allelic counts for human heart RNA-Seq samples.

## Supplementary Data

### Supplementary Figures

**Supplementary Figure S1.**
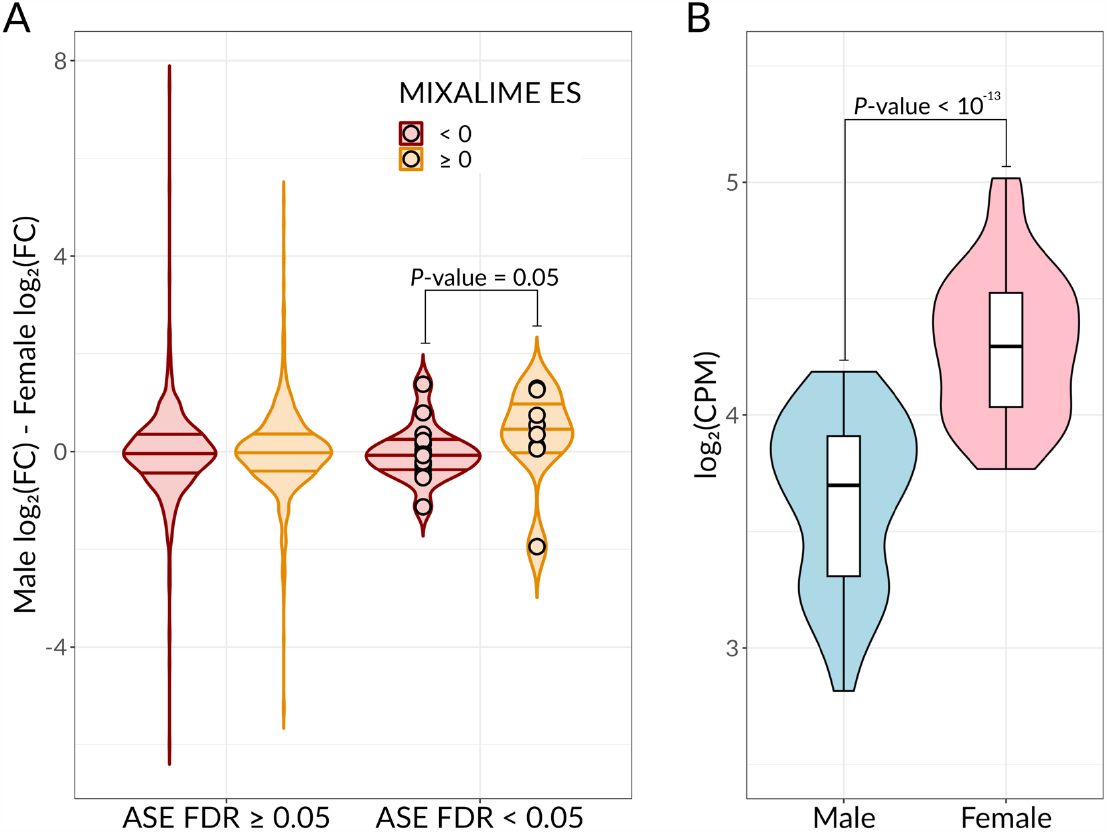
Verifying sex-specific ASEs called by MIXALIME. **A:** Y-axis: the relative difference between homozygous male and female samples, log_2_(Fold Change) of read counts between samples having both reference or both alternative alleles (0/0 and 1/1 for VCF GT field). X-axis: two SNP groups split by differential ASE FDR, ≥ or < 5%. Color denotes the ASE effect size, < or ≥ 0. *P*-value: Wilcoxon rank sum test. **B:** Violin plots representing the sex-specific expression of ZFX. Y-axis: log_2_(counts-per-million) estimated from gene counts of ^53^. X-axis: sex. *P*-value: Wilcoxon rank sum test. Abbreviations: CPM - Counts-Per-Million, ES - Effect Size, FC - Fold Change, FDR - False Discovery Rate.

**Supplementary Figure S2.**
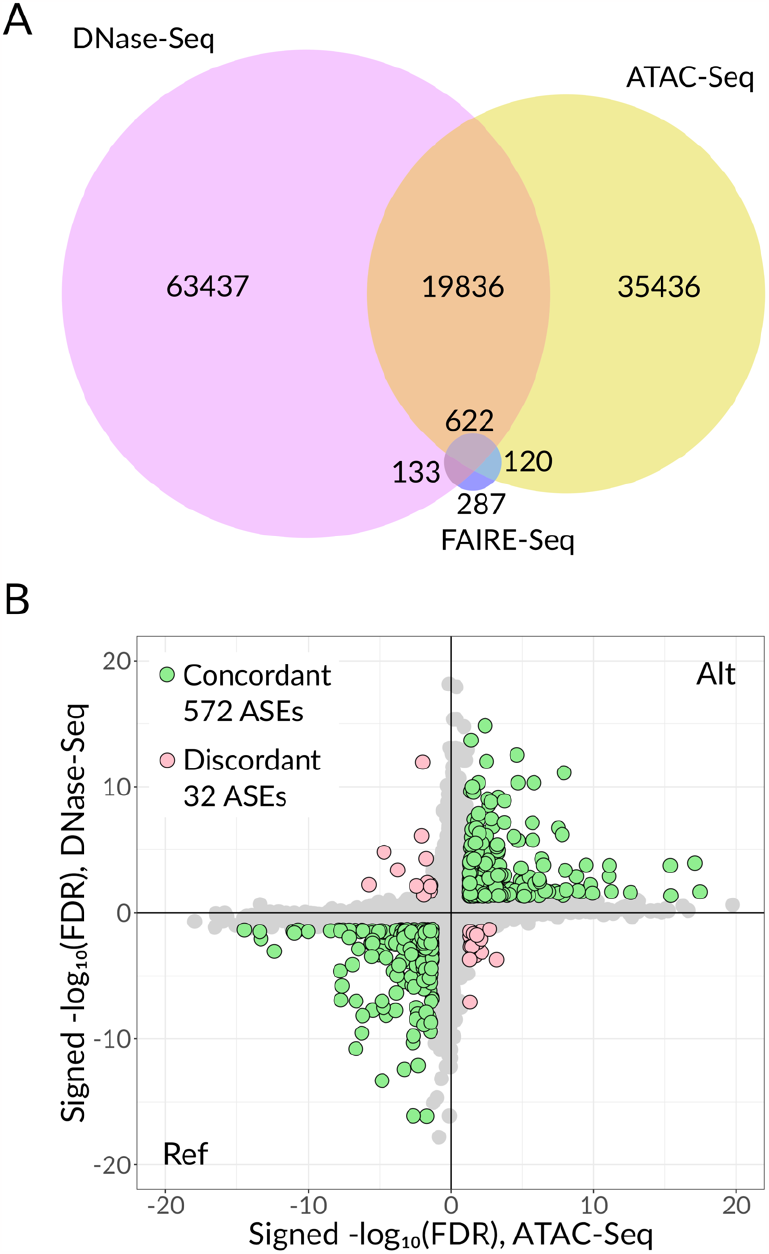
Comparison of UDACHA ASEs called in different experiment types. **A:** Venn diagrams for ASEs identified from DNase-Seq, ATAC-Seq, and FAIRE-Seq, unique rsSNPs carrying significant ASEs are counted. **B:** Scatter plot of ASEs significance from ATAC-Seq and DNase-Seq. X-axis: -log_10_(FDR) estimated from ATAC-Seq, the sign corresponds to the preferred allele, Ref < 0 and Alt > 0. Y-axis: signed -log_10_(FDR) from DNase-Seq. The color represents significance and concordancy (green: concordant, pink: discordant), and the gray dots denote SNPs with FDR > 5%. Only ASEs with significant allelic bias in both ATAC-Seq and DNase-Seq in the same cell type are checked for being concordant or discordant. Abbreviations: ASE - Allele-Specific Event, FDR - False Discovery Rate.

**Supplementary Figure S3.**
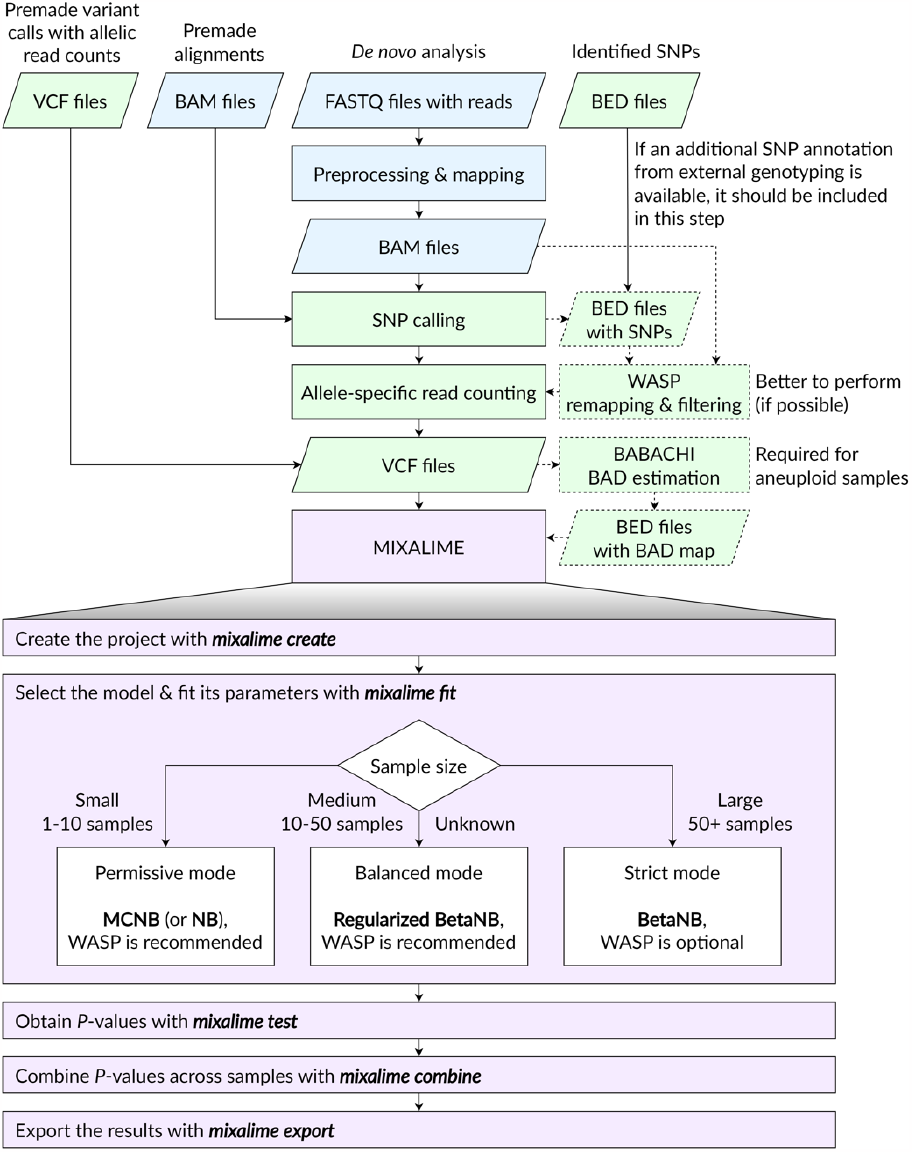
Schematic guidelines for MIXALIME practical application. MIXALIME allows direct ASE estimation from the VCF files generated by an SNP caller from BAM alignments. To obtain the input files for MIXALIME, the users may start from FASTQ files, align them to the genome, and perform SNP calling or, alternatively, use premade alignment files to call SNPs or the existing premade VCF files. In former cases, the WASP remapping and filtering procedure may be additionally performed to improve the ASE calling, especially for low sample sizes. If the external genotyping information is available it can be used to expand or filter the SNP calls. With the VCF files, BABACHI can be used to reconstruct the BAD maps for aneuploid samples before running MIXALIME. Finally, ASEs can be estimated with the MIXALIME selecting the model according to the sample size (see also main **Figure 4A**). Given the genotype calls are reliable (e.g. available from an external data or called from a large collection of samples), MIXALIME default filter requiring at least 5 reads at both alleles can be relaxed or omitted. Abbreviations: ASE - Allele-Specific Event, BAD - Background Allelic Dosage, BetaNB - Beta Negative Binomial, MCNB - Marginalized Compound Negative Binomial, NB - Negative Binomial, SNP - Single Nucleotide Polymorphism.

**Supplementary Figure S4.**
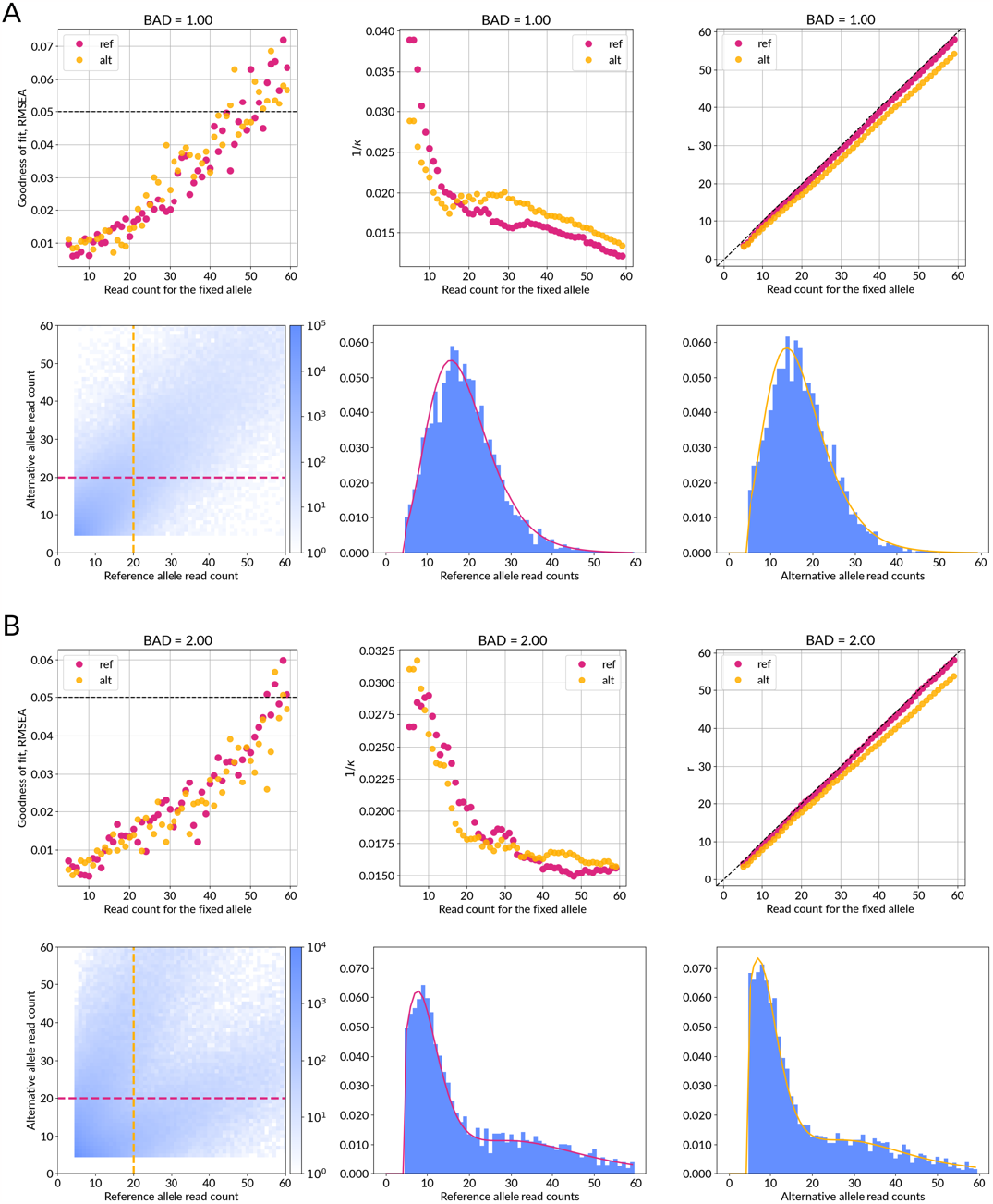
Visualization of the BetaNB model fits with MIXALIME. **A:** BAD = 1, **B:** BAD = 2. Both plots use the data from 98 DNase-Seq samples of **Figure 4B**. Parameter estimates were obtained with local local maximum likelihood. *r* is inferred from the line equation *r*(*z, a, b)* with *z* being the current sliding window position. Instead of concentration κ, dispersion 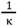 is plotted for visual clarity as κ tends to grow to infinity as coverage increases. Goodness of fit (RMSEA) was computed for particular slices corresponding to the window position. PDFs of those slices are drawn alongside heatmaps with the first figure being the PDF of the reference allele read count conditioned on the alternative allele read count and the second is vice-versa. Abbreviations: BAD - Background Allelic Dosage; RMSEA - Root Mean Square Error of Approximation

## Supplementary Methods

Detailed description of the MIXALIME statistical framework.

## Supplementary Tables

**Supplementary Table S1**. ASEs called by different MIXALIME models, QuASAR, and BaalChIP, and the intersection between ASEs and GTEx eQTLs / ADASTRA ASBs.

**Supplementary Table S2**. The results of Fisher’s exact test for association between eQTLs and ASEs called by different MIXALIME models with or without WASP pre-filtering.

**Supplementary Table S3**. The datasets used to evaluate the effect of BABACHI-generated BAD maps on the MIXALIME ASE calls.

**Supplementary Table S4**. The results of Fisher’s exact test for association between ASBs and ASEs called by MIXALIME basic and mixture models.

**Supplementary Table S5**. Sex-specific differential ASEs detected in heart CAGE data.

**Supplementary Table S6**. Overview of the datasets used to construct the UDACHA database.

**Supplementary Table S7**. Cell type-specific comparison of UDACHA ASEs and ADASTRA ASBs.

